# Engineering cardiolipin binding to an artificial membrane protein reveals determinants for lipid-mediated stabilization

**DOI:** 10.1101/2024.05.27.592301

**Authors:** Mia L. Abramsson, Robin A. Corey, Jan Škerle, Louise J. Persson, Olivia Andén, Abraham O. Oluwole, Rebecca J. Howard, Erik Lindahl, Carol V. Robinson, Kvido Strisovsky, Erik G. Marklund, David Drew, Phillip J. Stansfeld, Michael Landreh

**Affiliations:** Department of Microbiology, Tumor and Cell Biology, Karolinska Institutet, 171 65 Solna, Sweden; School of Physiology, Pharmacology & Neuroscience, University of Bristol, BS8 1TD Bristol, UK; Department of Biochemistry and Biophysics, Stockholm University, 104 05 Stockholm, Sweden; Institute of Organic Chemistry and Biochemistry, Academy of Science of the Czech Republic, 160 00 Prague, Czech Republic; Department of Chemistry – BMC, Uppsala University, 751 23 Uppsala, Sweden; Department of Biochemistry and Biophysics, Science for Life Laboratory, Stockholm University, 171 65 Solna, Sweden; Department of Chemistry, University of Oxford, OX1 3QZ Oxford, UK; Kavli Institute for Nanoscience Discovery, University of Oxford, OX1 3QU Oxford, UK; Department of Applied Physics, Science for Life Laboratory, KTH Royal Institute of Technology, 171 65 Solna, Sweden; School of Life Sciences & Chemistry, University of Warwick, CV4 7AL Coventry, UK; Department for Cell and Molecular Biology, Uppsala University, 751 24 Uppsala, Sweden

**Author notes:** Robin A. Corey, Philip J. Stansfeld & Michael Landreh, **E-mail:**, or.

## Abstract

Integral membrane proteins carry out essential functions in the cell, and their activities are often modulated by specific protein-lipid interactions in the membrane. Here, we elucidate the intricate role of cardiolipin (CDL), a regulatory lipid, as a stabilizer of membrane proteins and their complexes. Using the *in silico*-designed model protein TMHC4_R (ROCKET) as a scaffold, we employ a combination of molecular dynamics simulations and native mass spectrometry to explore the protein features that facilitate preferential lipid interactions and mediate stabilization. We find that the spatial arrangement of positively charged residues as well as local conformational flexibility are factors that distinguish stabilizing from non-stabilizing CDL interactions. However, we also find that even in this controlled, artificial system, a clear-cut distinction between binding and stabilization is difficult to attain, revealing that overlapping lipid contacts can partially compensate for the effects of binding site mutations. Extending our insights to naturally occurring proteins, we identify a stabilizing CDL site within the *E. coli* rhomboid intramembrane protease GlpG and uncover its regulatory influence on enzyme substrate preference. In this work, we establish a framework for engineering functional lipid interactions, paving the way for the design of proteins with membrane-specific properties or functions.

## Introduction

Biological membranes, which are vital for cellular life, provide a specific and highly adaptable lipid environment for membrane proteins that govern numerous cellular functions (*1*). The exact roles that the different membrane lipids play in the regulation of membrane proteins often go unacknowledged, as their highly dynamic interactions challenge conventional analytical methods. Despite these obstacles, evidence has consistently highlighted the crucial role of lipids (*2*), for example as allosteric regulators (*3*), facilitating protein oligomerization (*4*), or locally affecting the properties of the membrane (*5*). The simplest form of lipid-mediated regulation is the stabilization of specific protein conformations (*6*), resulting in the observation of individual lipid molecules in high-resolution structures (*7*). These “structural” lipids often display increased residence times at their binding sites which distinguish them from non-regulatory, “annular” lipids (*8*).

Cardiolipin (CDL) is a prime example of a lipid with regulatory activity for both bacterial and mitochondrial membrane proteins (*9*). Due to its unique structure, comprised of two phosphate groups which both potentially carry a negative charge, and four acyl chains, CDL mediates the assembly of membrane protein oligomers, for example in the respiratory chain supercomplexes (*10*). The double phosphate groups can create strongly attractive electrostatic interactions with basic side chains, which makes CDL an idea model lipid to understand interactions, but it also exhibits more specific patterns. Of note, both the head groups and all four acyl chains are thought to be important components of supercomplex stabilization (*11*). Similarly, CDL plays an essential role in the dimerization of the Na^+^/H^+^ antiporter NhaA, which increases the exchanger activity to protect the bacteria from osmotic stress (*12, 13*). In addition, CDL can affect the activity of other membrane proteins such as ADP/ATP carrier Aac2 and magnesium transporter MgtA by acting as an allosteric regulator (*14, 15*). Therefore, sites displaying preferential CDL binding may indicate lipid-activated regulatory mechanisms. To address this possibility, we have previously used coarse-grained molecular dynamics (CG-MD) simulations to map CDL binding sites on *E. coli* inner membrane proteins with published structures, identifying specific amino acids and binding site geometries that mediate preferential interactions with CDL (*16*). Although such CDL “fingerprints” are found in a wide range of proteins with different activities, they stop short of clarifying the functional role of lipids at these sites, with predictions of their functionality remaining largely speculative. Addressing this knowledge gap requires monitoring both the molecular interactions as well as the structure or stability of membrane protein complexes. For instance, thermal-shift assays provide data on lipid binding and associated changes in protein stability, which may indicate a functionally or structurally important lipid interaction (*17*). Moreover, native mass spectrometry (nMS) has gained traction for membrane protein analysis, revealing the influence of lipids on oligomerization (*18*), binding affinities (*19*), and conformational stability (*20*). Monitoring mass shifts captures individual lipid interactions across multiple protein populations, while gas-phase dissociation provides insight into lipid stabilization. nMS thus captures key features of regulatory lipid interactions, and is especially powerful when coupled with MD which provides insight at the atomistic level (*21*).

Being able to connect individual lipid binding events to the stability of a protein complex is a crucial step towards predicting functionally important CDL interactions. We reasoned that a combined MD and nMS strategy may reveal basic requirements for CDL-mediated stabilization. However, the sequence and structures of membrane proteins are evolutionarily entrenched with the lipid composition of their surrounding membrane. To reduce the system to first principles, we turned to TransMembrane Helical Core Tetramer_Rocket-shaped (TMHC4_R, hereafter referred to as ROCKET), an artificial membrane protein tetramer whose sequence was derived from Rosetta Monte Carlo calculations (*22*). ROCKET includes a generic lipid-water interface composed of a ring of aromatic residues and a ring of positively charged residues on the cytoplasmic side. Into the ROCKET scaffold, we designed several CDL binding sites based on our observations from *E. coli* proteins and tested their effect on tetramer stability using nMS. We find that local dynamics and the spatial distribution of charged residues distinguish stabilizing from non-stabilizing sites. However, we also observe that predicting the impact of individual mutations on lipid binding and stabilization from the structure can be challenging, even in our highly artificial system. These difficulties arise from the fact that lipid interactions are heterogeneous, and the loss of one type of contact may be compensated by another. Screening our database of *E. coli* CDL binding sites (https://osf.io/gftqa/) for binding sites that resemble stabilizing sites in ROCKET, we uncover a highly stabilizing CDL interaction in the membrane protease GlpG, which regulates the substrate preference of the enzyme. In summary, our study demonstrates the potential as well as the challenges in designing functional CDL sites on artificial proteins that can recognize membrane compositions.

## Results

### Design of a CDL binding site in ROCKET

As first step, we characterized the inherent lipid binding properties of ROCKET (Fig 1a) through CG-MD simulations of the protein in a mimetic *E. coli* membrane. The membrane composition was modeled with a distribution of POPE, POPG, and CDL in a ratio of 67:23:10, and the system was simulated for 5 x 10 µs while monitoring the lipid interactions. We observed abundant lipid interactions, with CDL displaying markedly more localized binding than POPE or POPG (Fig S1). The N-terminal region on the first transmembrane helix bound CDL with average occupancy of 71 % and average residence time of 35 ns (at R9). These values are extracted from the full 50 µs of simulation data. The site, which we termed Site 1, consists of three basic residues (R9, K10, and R13) and an aromatic residue (W12), which corresponds to a consensus CDL binding motif (Fig 1b) (*16*). We also observed a second site, involving W12 in a slightly rotated conformation, and R66 on helix 2 of the neighboring subunit. This site, termed Site 2, exhibited significantly lower occupancy of 56 % and an average residence time of 35 ns (at R66). W12 can engage in CDL binding at either site, including simultaneously both sites (Movie S1). Both sites represent distinct lipid binding modes: Site 1 is a high-occupancy site away from the protein core with extensive head-group interactions, and Site 2 is a lower-occupancy site with extensive acyl chain contacts close to the protein core. Note that, while the occupancies are high, the residence times are relatively low, as CDL is readily exchanged between the two sites. We decided to use these two sites, which arose from purely statistical distribution of charged and aromatic residues, as basis for engineering a stabilizing CDL site.

**Fig 1.**
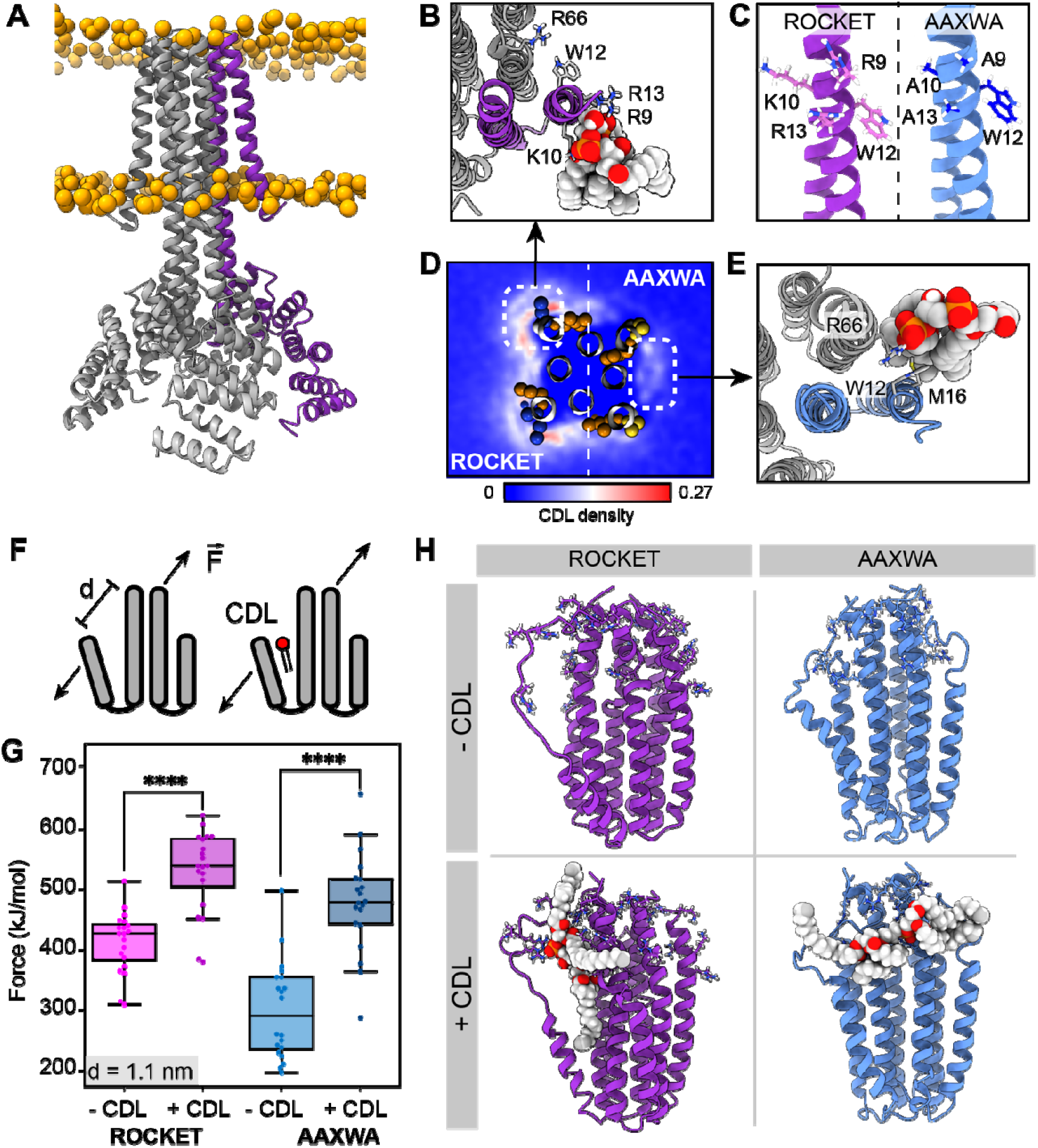
ROCKET contains two CDL binding sites with different structural implications. (A) Structure of ROCKET (PDB ID 6B85) in the membrane, with one protein subunit highlighted in purple and phosphate headgroups for the membrane shown as orange spheres. The structure was obtained from MemProtMD. (B) Top view of CDL binding to Site 1 taken from a 10 µs snapshot of a CG-MD simulation. The poses was converted to atomistic using CG2AT2 (23) to the CHARMM36m force field (24). CDL is shown as spacefill, and residues R9, K10, W12, R13, and R66 as sticks. The CDL-binding subunit is highlighted in purple. (C) Design of the ROCKET^AAXWA^ variant. Site 1 on helix 1 of ROCKET is shown on the left (purple), and ROCKET^AAXWA^ with the mutations R9A/K10A/R13A on the right (blue. (D) CG-MD-derived CDL densities around a heterotetramer composed of two ROCKET subunits (left) and two ROCKET^AAXWA^ subunits (right). Units are number density. Site 1 on ROCKET and Site 2 on ROCKET^AAXWA^ are highlighted by dashed boxes. R9, K10, W12, and R13 are shown as spheres (basic in blue, aromatic in orange). Densities are computed over 5 x 10 µs simulations. (E) Top view of CDL binding to Site 2 in ROCKET^AAXWA^ following CG-MD and converted to atomistic as per panel B. Interacting residues W12, M16, and R66 on the neighboring subunit (grey) are shown as sticks. (F) Setup of gas-phase MD simulations for unfolding of ROCKET and ROCKET^AAXWA^ with and without lipids. The placement of the lipid in the schematic is arbitrary. (G) Plots of the integral of the force required to separate helix 1 and 2 (d = 1.1 nm), for ROCKET (purple) (p=3.85*10-7) and ROCKET^AAXWA^ (blue) (p=2.93*10^-8^) with and without bound CDL show a more pronounced increase in stability for CDL-bound ROCKET^AAXWA^ compared to ROCKET (two-tailed t-test with n=20).(H) Snapshots from gas-phase MD simulations show broad interactions of CDL across the subunits of lipid-bound ROCKET^AAXWA^ (blue) and more localized interactions with fewer intermolecular contacts for ROCKET (purple). Amino acid position 9, 10, 12, 13, and 66 are shown as sticks in each subunit.

To separate the two CDL binding modes, we generated ROCKET mutants *in silico* and performed CG-MD with two subunits each of ROCKET and ROCKET mutants. We found that substituting the charged residues of Site 1 with alanine (R9A/K10A/R13A, Fig 1c) redirected preferential CDL binding to Site 2 (Fig 1d, e). In this mutant, which we termed ROCKET^AAXWA^, the Site 2 had an occupancy of 53 % and an increased average residence time of 47 ns (R66), whereas Site 1 had a reduced occupancy of 45 % and a residence time of 45 ns (R9A). This occupancy difference was quantified by CG simulations and showed significant reduction of total CDL binding between ROCKET and ROCKET^AAXWA^ (Fig 1d, S1).

Next, we evaluated the potential for lipid-mediated stabilization at both sites using gas-phase atomistic MD simulations, which allows for a direct comparison with nMS. We applied a pulling force between two adjacent subunits of ROCKET and ROCKET^AAXWA^ tetramers with and without bound CDL and determined the force required to separate the protein chains (Fig 1f). Analysis of the secondary structure content shows that the AAXWA mutation stabilizes the conformation of helix 1 (Fig S2). However, we found that separating the adjacent helices from the neighboring subunits of ROCKET^AAXWA^ by 1.1 nm, the point at which non-covalent interactions between the transmembrane helices are disrupted, required more force when CDL was present (Fig 1g, Fig S2). ROCKET, on the other hand, displayed a lower significant difference in force with or without CDL. Snapshots from the simulations reveal that the lipid forms multiple contacts with both subunits adjacent to Site 2 in ROCKET^AAXWA^, which likely gives rise to the stabilizing effect (Fig 1h). We conclude that channeling the CDL molecules to inter-helix sites may be a prerequisite for lipid-mediated stabilization.

### Inter-helix CDL binding stabilizes ROCKET^AAXWA^ in the gas-phase

Having derived two ROCKET variants with distinct CDL binding modes from MD simulations, we turned to cryogenic electron microscopy (cryo-EM) and nMS to investigate their lipid interactions experimentally. We first analyzed ROCKET and ROCKET^AAXWA^ in the presence of CDL by cryo-EM. The resulting density maps for each protein with no other particle class detected, refined to a resolution of 3.8 and 3.9 Å respectively, show essentially identical architectures that agree with the previously solved crystal structure of ROCKET (*22*), confirming that the mutations do not disrupt the native structure (Fig S3, Table S1). Although a definitive atomic-level molecular model was not possible at this resolution, we also observed in both maps a diffuse non-protein density which partially overlaps the head-group of CDL predicted in Site 2 (Fig S3). Interestingly, we see no extra density in Site 1, however, this site is more exposed, making it more likely that excess detergent can outcompete the binding of CDL in this site. Furthermore, the orientation of CDL is more flexible in Site 1 than Site 2 (Figure 1D), which also reduces the likelihood of obtaining a sufficiently defined density. To determine lipid binding preferences, we therefore reconstituted the proteins into liposomes composed of polar *E*.*coli* polar lipid extracts (Fig 2a). By releasing the proteins from the liposomes inside the mass spectrometer and monitoring the intensity peaks corresponding to apo- and lipid-bound protein, we can compare lipid preferences of both variants (Fig 2a). We find that tetrameric ROCKET retains up to three CDL molecules, which can be identified by their characteristic 1.4 kDa mass shift, as well as a significant number of phospholipids between 700 and 800 Da (Fig 2b). The data thus show a preference for CDL, which constitutes only 10% of the liposome. Interestingly, nMS of ROCKET^AAXWA^ revealed a similarly specific retention of up to three CDL molecules for 17+ charge state, although the intensity of the lipid adducts was reduced by approximately 50% (Fig 2c). The mass spectra show that the preference for CDL is preserved in the ROCKET^AAXWA^ variant, while the occupancy is reduced, indicating either lower affinity in solution or lower stability of the protein-lipid complex in the gasphase.

**Fig 2.**
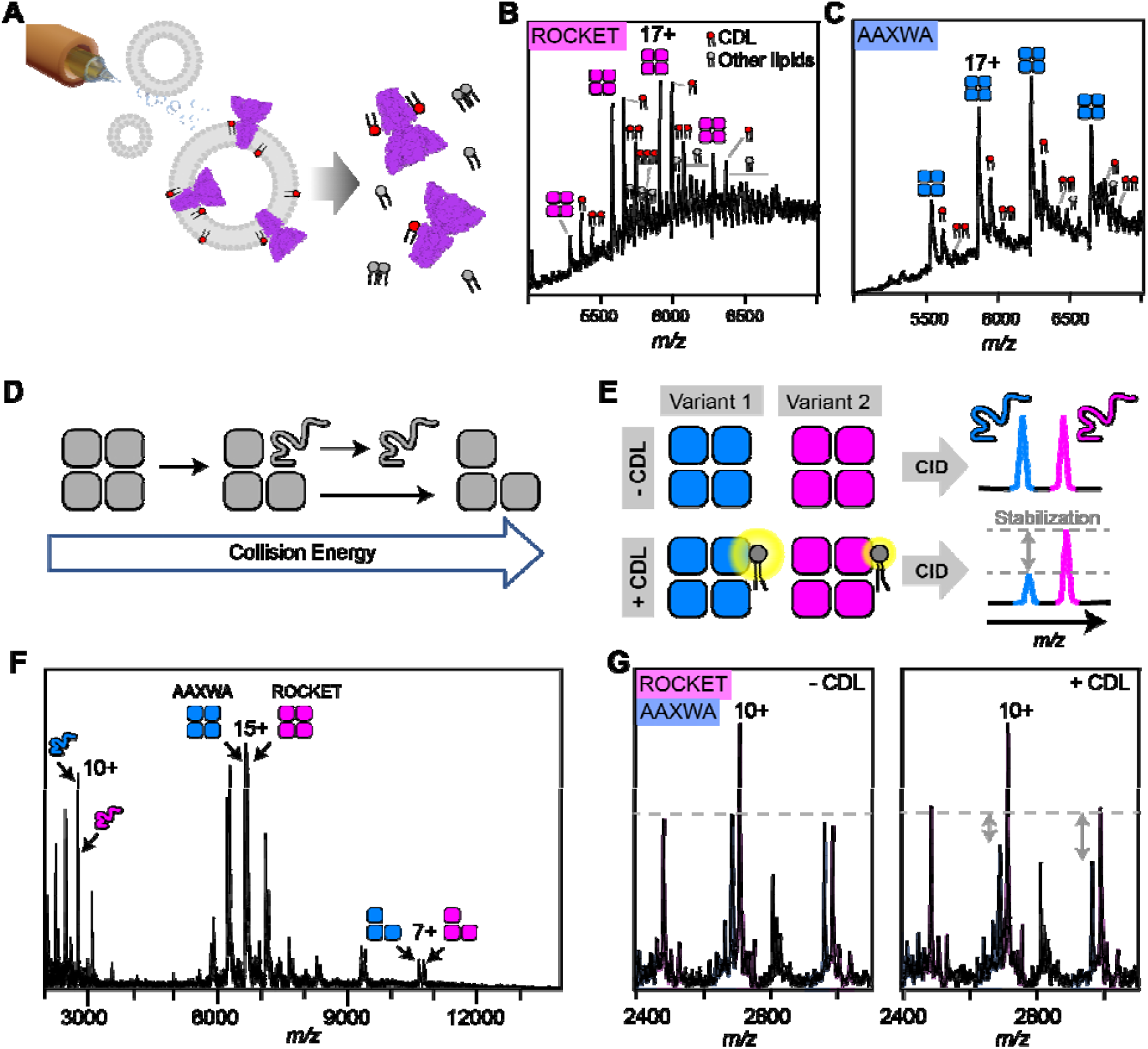
nMS analysis of lipid binding and lipid mediated stabilization of ROCKET and ROCKET^AAXWA^. (A) Schematic depiction of electrospray ionization (ESI) process for proteo-liposomes, leading to the ejection of protein-lipid complexes into the gas-phase. (B) A representative mass spectrum of ROCKET released from proteoliposomes shows tetramers with 1-3 bound CDL molecules, as judged by the characteristic mass shift of 1.4 kDa, as well as additional lipids with molecular weights between 700 and 800 Da. (C) Release of ROCKET^AAXWA^ from proteoliposomes shows retention of CDL molecules. For the 18+ ion of the tetramer, a maximum of three CDL adducts can be assigned unambiguously. The reduced lipid adduct intensity compared to ROCKET indicates reduced lipid binding and/or complex stability. (D) Schematic illustrating the process of gas-phase subunit unfolding and ejection from ROCKET tetramers at increasing collision energies. (E) A nMS assay to assess CDL-mediated stabilization of ROCKET and ROCKET^AAXWA^. Simultaneous dissociation of ROCKET and ROCKET^AAXWA^ leads to the ejection of unfolded monomers, which can be quantified by nMS (top row). Addition of CDL to the same mixture results in lipid binding to tetramers. If CDL binding stabilizes one tetrameric variant more than the other, the amount of ejected monomers will be reduced accordingly (bottom row). (F) Representative mass spectrum of ROCKET and ROCKET^AAXWA^ at a collision voltage of 220 V. Intact tetramers are seen in the middle, ejected monomers and stripped trimers are seen in the low and high m/z regions, respectively. (G) Zoom of the low m/z region of a mixture of 25 µM each of ROCKET and ROCKET^AAXWA^ with a collision voltage of 200 V before (left) and after (right) the addition of 50 µM CDL. The three main charge states for both variants can be distinguished based on their mass difference. Addition of CDL reduces the intensity of the ROCKET^AAXWA^ monomer peaks compared to ROCKET (dashed line).

To validate these findings, we analyzed lipid binding to detergent-solubilized ROCKET. First, we optimized detergent conditions for maintaining the intact ROCKET tetramer (Fig S4). Importantly, we observe no co-purified lipids, indicating that detergent can remove CDL during extraction, as expected for lipids with short residence times.(*21*) We then performed a competition assay where we mixed both variants in C8E4 detergent containing a limiting amount of CDL (50 µM) and monitored binding with nMS. As expected, both variants bound three distinguishable CDL molecules per tetramer, however, ROCKET displayed significantly more intense lipid adducts than ROCKET^AAXWA^ (Fig S4). In CG-MD simulations, the overall CDL occupancy is lower in ROCKET^AAXWA^ than in ROCKET, meaning fewer lipids will be bound simultaneously. The nMS data show CDL retention by both variants, but the ROCKET^AAXWA^ protein has lower-intensity CDL adduct peaks (Figure 2B, C). This finding suggests that both variants bind CDL, but in the ROCKET^AAXWA^ variant, the sites have lower occupancy. The nMS data are therefore consistent with CDL binding preferentially to Site 1 in ROCKET and preferentially to Site 2 in the ROCKET^AAXWA^ variant.

Next, we explored how CDL binding to either site affects the stability of ROCKET, using the oligomeric state in nMS as a measure. To avoid interference from different lipids in the reconstituted liposome system, we switched to detergent micelles as vehicles for nMS and employed gas-phase dissociation of the intact protein complexes to remove bound detergent. Briefly, collisions with gas molecules in the ion trap of the mass spectrometer cause thermal unfolding of a single subunit in the complex, which is then ejected as a highly charged, unfolded monomer (Fig 2d) (*25*). By comparing the peak intensities of the monomers that are ejected simultaneously from two protein oligomers, we can obtain information about their relative stabilities. Therefore, by adding CDL to an equimolar mixture of ROCKET and ROCKET^AAXWA^, dissociating the resulting complexes, and monitoring changes in monomer signal intensities, we can determine whether lipid binding to Site 1 or Site 2 affects tetramer stability (Fig 2e). Importantly, by comparing changes in peak intensities with and without CDL while keeping all other conditions constant, we can avoid interference from changes in gas-phase fragmentation or ionization efficiency. nMS of ROCKET and ROCKET^AAXWA^ shows the release of highly charged monomers which can be distinguished based on their masses (Fig 2f). We then added CDL to the protein solution and repeated the measurement using identical conditions. We observed a reduction in the peak intensities of ROCKET^AAXWA^ monomers compared to ROCKET (Fig 2g). We do not observe a change in the charge state distributions for tetramers or monomers, or notable fragmentation. Therefore, the change in monomer ratio suggests that CDL stabilizes the ROCKET^AAXWA^ tetramer to a greater extent than the ROCKET tetramer. These findings are surprising, since the AAXWA variant displays significantly lower lipid binding (Fig 2 B, C). However, considering the predictions from CG- and gas-phase MD, the increase in stability can be attributed to the preferential binding of CDL to the inter-helix Site 2 in the AAWXA variant. CDL binding to the distal Site 1, as preferred in ROCKET, involves fewer intermolecular contacts, and is therefore unlikely to exhibit a similarly stabilizing effect.

### Multiple structural features impact CDL-mediated stabilization

The finding that inter-helix CDL binding stabilizes a tetrameric membrane protein in the gas-phase recapitulates a key feature of both prokaryotic and eukaryotic membrane proteins (*18, 26*). Unlike naturally evolved proteins, however, the extraordinary stability and mutation tolerance of the ROCKET scaffold enables us to dissect further the requirements for CDL-mediated stabilization. We therefore applied the above MS strategy to quantitatively assess lipid-mediated stabilization in our model system between two protein variants using ROCKET^AAXWA^ as an internal reference. We can determine relative stability changes upon CDL addition for different ROCKET variants by plotting the ratios of the total intensity of the peaks for monomeric ROCKET mutants (ROCKET^MUT^) to the total intensity of all protein monomer peaks in the spectrum (ROCKET^MUT^ + ROCKET^AAXWA^) with and without CDL. If ROCKET^AAXWA^ is stabilized more than the variant of interest, the ratio increases with CDL addition (Fig 3a). As expected, the AAXWA mutation significantly increased the stabilizing effect of CDL, as determined from four independent repeats (Fig 2g, 3b, and 3c). With this assay, we then explored whether introducing different structural features into Site 1 could turn it into a stabilizing CDL binding comparable to Site 2. As a first hypothesis, we reasoned that a destabilization of the core of ROCKET might increase the effect of CDL. We introduced a destabilizing mutation (A61P) in helix 2, right below the headgroup region, theorizing that the proline-induced kink would destabilize the ROCKET tetramer. AlphaFold2 predictions (*27, 28*) indicated that the ROCKET^A61P^ mutation does not affect the tetrameric state (*29*), which was confirmed by nMS. To our surprise, ROCKET^A61P^ exhibited significantly less CDL stabilization than ROCKET^AAXWA^, and was comparable to ROCKET (Fig 3d, Fig S5). This observation indicates that the introduction of a proline in the protein core does not sufficiently destabilize the protein to be counteracted by lipid binding.

**Fig 3.**
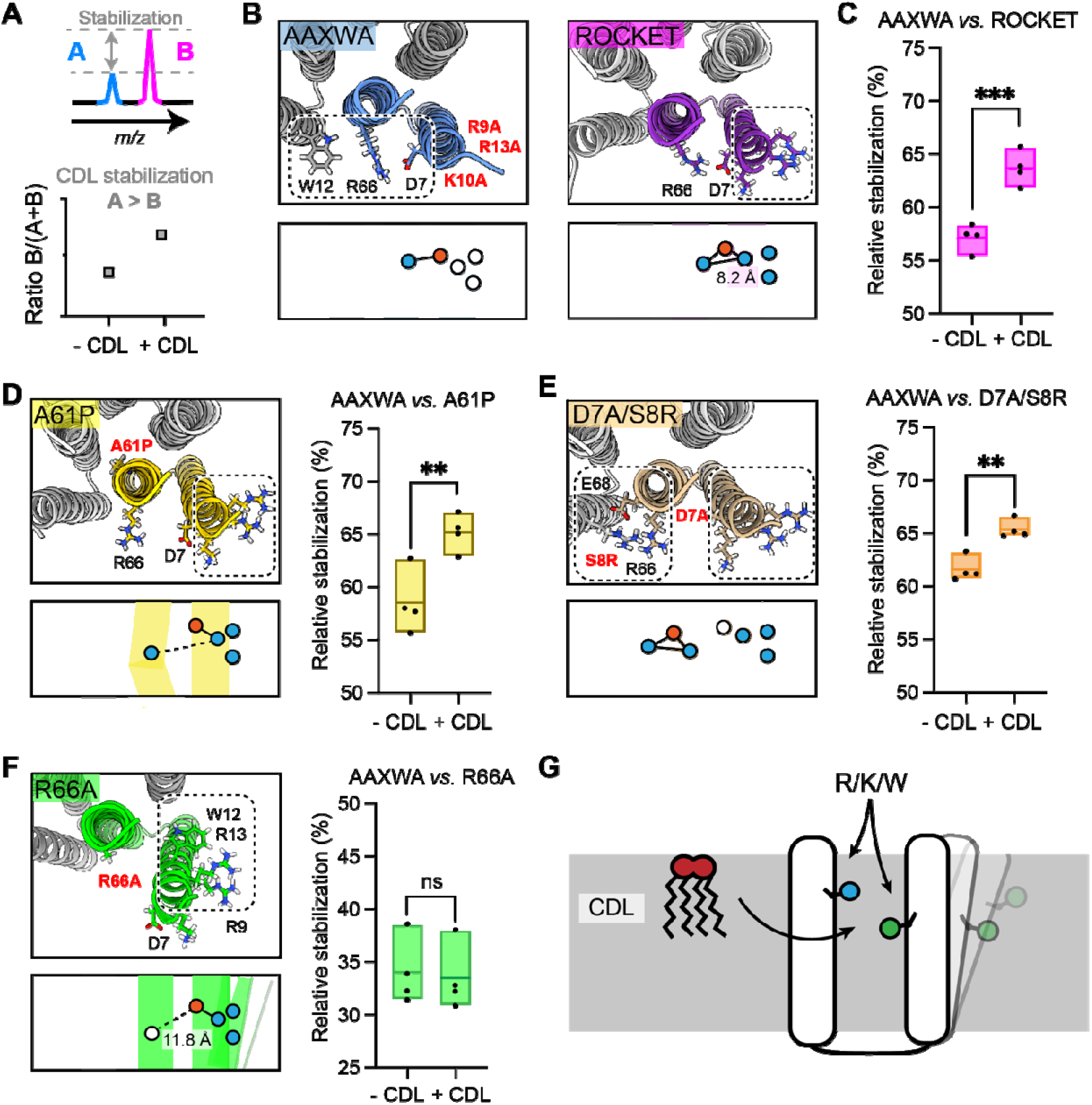
Assessment of lipid-mediated stabilization effects on ROCKET variant. (A) Principle for pairwise analysis of protein stabilization by CDL. Peaks representing monomers released of protein A and B display intensity changes upon lipid addition. Plotting the peak intensities as a ratio of B to total protein (A + B) shows an increase upon CDL addition, if A is stabilized more than B. (B) Residues involved in CDL headgroup binding to ROCKET and ROCKET^AAXWA^ were derived from CG-MD simulations (Fig 1) and are shown based on AlphaFold2 models as top view, with the area occupied by CDL as a dashed rectangle. A single subunit is colored (purple for ROCKET and blue for ROCKET^AAXWA^). Below the structure, the orientation of the CDL site and the helices are shown as schematics. Positively and negatively charged residues are blue dots, respectively. Residues mutated to alanine are shown as white dots. (C) Plotting the peak intensity ratios of ROCKET to (ROCKET^AAXWA^ + ROCKET) in the presence and absence of CDL shows a decrease in ROCKET^AAXWA^ monomers when CDL is added (p=0.0007, two-tailed t-tests with n=4). (D) The CDL binding site and location of the A61P mutation (yellow) mapped on the AlphaFold2 model of ROCKET^A61P^ and shown as a schematic as a side view below. Intensity ratios show significantly more pronounced stabilization of ROCKET^AAXWA^ than ROCKET^A61P^ (p=0.0084, two-tailed t-tests with n=4). (E) Introduction of a second CDL binding site in the D7A/S8R variant (orange) mapped on the AlphaFold2 model of ROCKET^D7A/S8R^ and shown as a schematic as a side view below. The shift in intensity ratios show that ROCKET^AAXWA^ is still stabilized to a greater extent (p=0.0019, two-tailed t-tests with n=4), but with a smaller margin than ROCKET or ROCKET^A61P^. (F) The R66A mutation, designed to disconnect helix 1 from the tetrameric protein core, results in an outward rotation of the CDL binding site, as shown in the AlphaFold2 model (green) and the side view schematic. Intensity ratios show no change upon CDL addition, suggesting that ROCKET^R66A^ is stabilized to a similar extent as ROCKET^AAXWA^ (p=0.8113, two-tailed t-tests with n=4).(G) Conceptual diagram depicting structural features that promote CDL-mediated stabilization. Distributing the residues that interact with the lipid headgroup, usually basic and aromatic residues, between two helices, as well as Involvement of flexible protein segments, indicated by an outward movement of the right helix, also enhances stabilization by CDL.

As second hypothesis, we reasoned that additional lipid binding sites may increase the effect of CDL binding (Fig S6). We therefore mutated residue D7 to alanine and S8 to arginine. The D7A/S8R variant retains the high-affinity Site 1 on helix 1, but includes an additional site composed of R8 on helix 1 and R66 and E68 on helix 2 which does not overlap with Site 1 (Fig 3d). Quantification of the monomer release with and without CDL suggests a shift towards increased stability with CDL, albeit not as pronounced as for ROCKET^AAXWA^ (Fig 3d, Fig S5). This data leads us to speculate that Site 1 may still bind the bulk of the available CDL molecules, or that the salt bridge S8-E68 may contribute increased stability in a CDL-independent manner.

Having explored directed lipid binding (AAXWA), increased lipid binding (D7A/S8R) and core destabilization (A61P), we reasoned that the flexibility of the CDL binding site may affect stabilization. This feature is challenging to implement in the ROCKET scaffold, since it is designed around a tightly folded hydrogen bond network with a melting temperature of > 90°C (*22*). We therefore decided to untether helix 1 from the core of the protein by mutating R66, which forms a salt bridge with D7, to alanine. The AlphaFold2 model shows helix 1 being tilted away form the core, creating a large hydrophobic gap in the transmebrane region and turning Site 1 towards the neighboring subunit (Fig 3f, Fig S5). Interestingly, ROCKET^R66A^ showed lower signal intensities in native mass spectra than all other variants, which may indicate overall lower stability in detergent. Quantification of the monomer release with and without CDL revealed no significant difference compared to ROCKET^AAXWA^, which means CDL binding has a stabilizing effect on both proteins (Fig 3f). This finding is surprising, since loss of R66 should result in increased binding to Site 1 and thus not stabilize the protein. However, untethering helix 1 may create an opportunity for CDL coordinated by W12 and/or R9-R13 to insert its acyl chains into the resulting inter-helix gap. In this manner, CDL could exert a stabilizing effect in the absence of preferential headgroup interactions.

From the designed ROCKET variants, we can conclude that structure-based predictions of stabilizing CDL interactions is challenging, as they arise from a combination of headgroup- and acyl chain interactions, as well as from their impact on the local structural dynamics of the protein. However, from our observations, we can conclude that CDL binding involving different helices, as in ROCKET^AAXWA^, and connecting flexible regions, as in ROCKET^R66A^, gives rise to the most pronounced CDL stabilization of our system (Fig 3g). Core destabilization, as well as introduction of additional headgroup contacts, had less of an impact, although the specific properties of the engineered protein scaffold may mitigate potential effects to some extent.

### Identification of a stabilizing CDL binding site in the *E*.*coli* rhomboid intramembrane protease GlpG

As outlined above, the features that cause CDL-mediated stabilization of the ROCKET scaffold were designed based on observations from CG-MD investigation of CDL interactions with of *E. coli* membrane proteins. We therefore asked whether the same features could indicate stabilizing, and by extension, functionally relevant CDL interactions in naturally occurring proteins. To test this hypothesis, we evaluated our database of monomeric *E. coli* membrane proteins CDL sites (https://osf.io/gftqa/) for the sequence distribution of basic residues to find binding sites that span multiple helices. We reasoned that if two or more basic residues that interact with the same CDL molecule are located further apart in the sequence than approximately 30 positions, they have a high likelihood of being on separate helices, whereas sites spanning less than 10 residues are confined to a single helix or loop. Plotting the maximum sequence distance on a log scale reveals a bimodal distribution, with 75 CDL sites spanning less than 30 positions, and 180 sites spanning more than 30 positions (Fig 4a, Fig S5). Site 1 in ROCKET is in the first group, with four positions between R9 and R13. This approach does not consider interfacial CDL molecules in homo-oligomers, which may bind *via* single residues on different subunits. We therefore limited the dataset to monomeric proteins with CDL sites spanning >30 residues which we manually inspected to find CDL sites linking potentially flexible regions.

**Fig 4.**
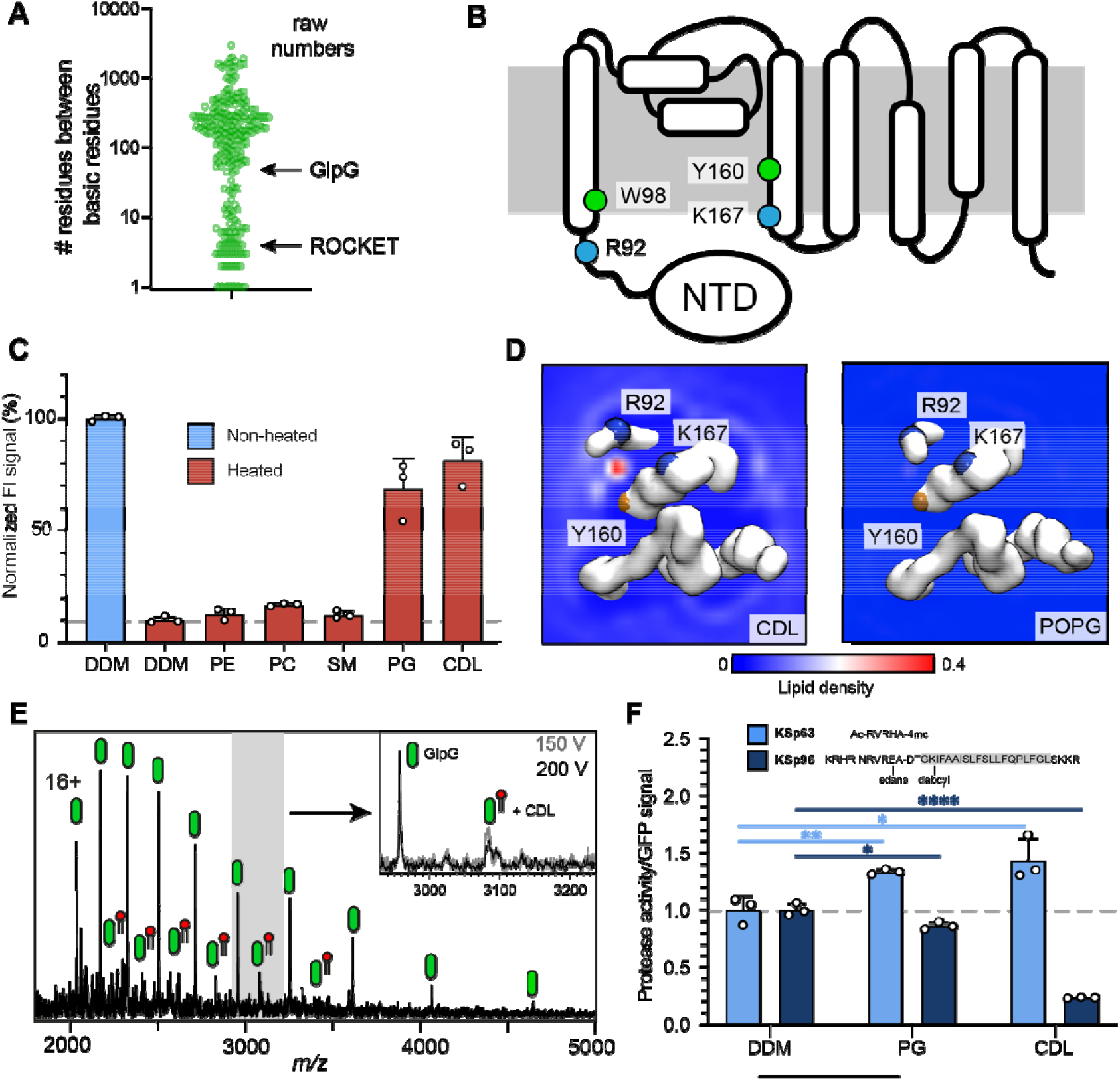
Identification of a functional cdl binding site in the glpg membrane protease. (a) distances between basic residues within the same cdl binding site and their occurrence in e. Coli membrane proteins. Glpg contains a cdl binding site with two basic residues separated by 75 positions. Site 1 in rocket (four positions) is indicated for reference. (b) topology diagram of glpg indicating the locations of basic (r; arginine, l; lysine) and aromatic (w; tryptophan, y; tyrosine) residues that interact with cdl. (c) gfp-thermal shift assay of glpg in detergent or in the presence of pe, pc, sm, pg, or cdl. The average fluorescence intensity (fi) indicates the fraction of soluble protein before (unheated) or after heating to 63 °c and removing precipitated protein. Data are normalized against the non-heated control in detergent. All measurements were performed in triplicates (n = 3). (d) cg-md-derived lipid density plots for cdl (left) and popg (right) around the transmembrane region of glpg (pdb id 2ic8) viewed from the cytoplasmic side. Units are number density. Cdl, but not popg, exhibits preferential binding to the r92-k167 site. The backbones of r92, k67, and y160 are shown as spheres (basic in blue, aromatic in orange). (e) nms spectra of glpg released from detergent micelles show a 1.4 kda adduct which can be removed through collisional activation of the protein. (f) glpg proteolytic cleavage rates (specific activity, by normalizing to the gfp signal) for soluble (extramembrane) substrate (ksp63) or transmembrane substrate (ksp96, transmembrane helix shaded) in the presence of pg or cdl. Both lipids increase the cleavage rate for the soluble substrate by approx. 40%. For the transmembrane substrate, addition of pg causes a moderate reduction in cleavage activity, whereas cdl causes near-complete inhibition of cleavage (cdl ksp96 p<0.0001, cdl ksp63 p=0.028, pg ksp96 p=0.0171, and pg ksp63 p=0.0089, all analyzed with two-tailed t-test, n=3).

The rhomboid intramembrane protease GlpG from *E*.*coli* contains a CDL site between R92 on helix 1 and K167 on helix 4, as well as one aromatic residue on each (W98 and Y160). Importantly, R92 borders on a disordered linker region that connects helix 1 to an N-terminal cytoplasmic domain (Fig 4b). These features suggest that CDL binding may have structural implications. We therefore measured whether any lipids imparted thermal stabilization in a GFP-based thermal shift assay (*17*). Measuring the fraction of GFP-tagged soluble protein after heating to 63 °C in the presence of different lipids showed that phosphatidyl ethanolamine (PE), phosphatidyl choline (PC), as well as sphingomyelin (SM) (which is not found in *E. coli*) had no stabilizing effect, whereas phosphatidyl glycerol (PG) and CDL resulted in near-complete protection form heat-induced precipitation (Fig 4c).

When performing CG-MD of GlpG in the model *E. coli* bilayer, we observed that CDL bound exclusively to the predicted site between helix 1 and 4, with an occupancy of 96.2% with an average residence time of 170 ns, but no preferential POPG binding was observed anywhere on the protein (Fig 4d, Fig S7). On the cytoplasmic side, the non-conserved residues W136 and R137 can potentially interact with CDL, however no increased CDL density was observed (Figure S7). The data suggest that the stabilizing effect of POPG in the thermal shift assay is due to binding to this site in the absence of CDL. The near-100% occupancy by CDL in the mixed bilayer MD simulations suggests that CDL outcompetes POPG when both lipids are present. To test whether GlpG binds CDL in the membrane, we used nMS analysis. We have previously noted a 1.4 kDa adduct on purified GlpG (*30*) and monomeric GlpG with sufficient activation to remove DDM micelle. Collisional activation revealed preferential loss of the adduct as a single species, confirming that it is a single lipid molecule, *i*.*e*. CDL (Fig 4e). These findings are in good agreement with the presence of four lipid tails in total, which likely stem from a single CDL in this location in the crystal structure of GlpG (Fig S7), although these were tentatively modeled as a PA and a PE molecule next to each other in the structure (*31*). The apparent high specificity for CDL prompted us to investigate the effect of CDL and PG on GlpG protease activity. For this purpose, we used a cleavage assay with fluorescently labeled substrates that represent either a soluble “extramembrane” substrate (KSp63) or transmembrane helix (KSp96) peptides containing the same GlpG cleavage site (*32, 33*). We found that PG and CDL both have a positive effect on the cleavage of the soluble substrate KSp63 compared to detergent-only, with both lipids increasing the cleavage rates by approximately 40% (Fig 4f). PG mildly reduced the cleavage rate of the transmembrane substrate KSp96 to approximately 90% compared to detergent-only. Strikingly, CDL caused drastic inhibition of transmembrane substrate cleavage, with only 20% activity remaining (Fig 4f). To find out how the lipid binding site, which is located on the cytoplasmic side, can affect substrate access to the active site, which faces the periplasm, we performed CG-MD simulations of the open and closed conformations of GlpG (*34*). We found that both conformations bound CDL in the same place, which agrees with the observation that CDL affects the flexibility of the linker connecting the N-terminal domain (*34*). Interestingly, the acyl chains of the tightly bound lipid extend into the gap between helices 2 and 5 (Fig S7), the lateral gate through which transmembrane substrates access the active site of GlpG (*34*). Loop 5, whose movement is critical for catalytic activity, remains unaffected, providing a rationale for the selective inhibition of transmembrane substrate cleavage by CDL. Although the biological role of CDL-regulated substrate preferences of GlpG remains to be clarified, our results demonstrate the identification of a functionally relevant CDL binding site in an *E. coli* membrane protein based on the insights from artificial protein design.

## Discussion

In this study, we have systematically investigated the requirements for CDL binding and stabilization of membrane proteins, using the scaffold protein ROCKET as a model system. CDL binding sites have few sequence requirements, which is evident from the fact that ROCKET contains multiple interaction sites solely from statistical distribution of aromatic and charged amino acids at the membrane interface (Fig 1). These sites exhibit characteristic features, including electrostatic interactions with the negatively charged lipid headgroups and flanking aromatic residues that align the acyl chains with the hydrophobic transmembrane domain. By introducing mutations at these sites, we can probe their contribution to CDL-mediated stabilization with native MS. To our surprise, we find that although CDL binding sites share well-defined structural features, mutations of these features does not always yield predictable structural effects. Instead, their ability to contribute to lipid-mediated stabilization depends on multiple additional factors. For example, mutations that reduce CDL binding to a high-affinity site can cause redistribution to lower-affinity sites that have, in turn, a more pronounced effect on stability. This appears to be the case for *ROCKET*^*AAXWA*^. Furthermore, mutations that reduce lipid binding at a low-affinity site may change the local stability of the protein, and results in an increase in lipid-mediated stabilization, as seen in *ROCKET*^*R66A*^. Lipid interactions are dynamic and heterogeneous, and mutating a well-defined CDL site can give rise to multiple compensatory interactions which in turn affect the local protein environment.

Despite these challenges, the different CDL binding sites in ROCKET demonstrate that commonly observed structural features (the number of headgroup-interaction residues, local flexibility, and the involvement of multiple transmembrane helices) impact lipid-mediated stabilization to different extent. Building upon the insights garnered from ROCKET, we extended our investigation to *E. coli* proteins and identified GlpG as a case study for CDL’s impact on function. Previous studies have shown that GlpG conformation is impacted by the surrounding lipids and membrane geometry (*30, 31*), and CDL has been suggested to affect proteolytic activity (*35*). Our observation that CDL acts as an allosteric activator for the cleavage of soluble substrates while exerting an inhibitory effect on the processing of transmembrane substrates establishes CDL as a regulator of GlpG activity. Similarly, a CDL binding site (W70/R71/R151) in the yeast ATP/ADP carrier Aac2 has the same hallmarks as the GlpG site (Figure S8). CDL binding to this site connects the flexible N-terminal transmembrane helix to the protein core, resulting in stabilization of the tertiary structure and increased transport activity. nMS shows that the protein retains co-purified CDL.(*14*) These findings suggest that the features that mediate lipid stabilization in ROCKET are hallmarks of functionally important lipid binding sites in native membrane proteins.

Although we can identify relatively intuitive features of CDL interaction sites, we find that the connection between lipid binding and stabilization is not clear-cut. For example, one destabilizing mutation increases lipid-mediated stabilization whereas another does not (compare ROCKET^R66A^ and ROCKET^A61P^). Furthermore, being able to bind more lipids do not translate to forming more stable complexes (compare ROCKET and ROCKET^AAXWA^). The reasons are likely two-fold: Firstly, our approach has methodological limitations, as gas-phase stability is not easily correlated with condensed phase stability. In case of CDL, increasing the number of molecular contacts likely translates to stabilizing effects in both phases. However, charge interactions are relatively strengthened in the gas phase, whereas some hydrophobic contacts will be lost. For example, CDL sites contain aromatic residues (W or Y) close to the ester bonds of the lipids, which likely serve to orient the lipids, but the roles of which have not been examined in ROCKET. Unlike charge interactions with lipid head groups, such subtle contributions are likely distorted by the transfer to the gas phase, making it difficult to confidently assign changes in stability or lipid occupancy. Furthermore, the bulk of the nMS analysis is done using detergents, which have delipidating effects, meaning we will observe fewer lipid binding events and lower occupancy in nMS than in the CG-MD simulations.(*21*) Collisional activation required to remove the detergent micelle or lipid vesicle additionally strips away lipids that bind predominantly via hydrophobic interactions.

To avoid this limitation of our MS approach, we have focused here CDL bound via direct headgroup contacts, which can be readily predicted with CG-MD and analyzed with nMS. Secondly, our approach highlights that protein-lipid interactions are complex. The ROCKET scaffold is, *per* design, extremely stable and has no function. Thus, it does not capture the dynamic nature of most membrane proteins, which, in turn, is often related to lipid-mediated regulation (*12*). The key finding of our study is that emulating the architecture of a binding site is only the first step to uncovering the principles of lipid regulation, and that the effect of lipids on the local dynamics are critical to understanding how they shape membrane protein function. Integrated design approaches based not only on static structures but on dynamic models are therefore the key to designing membrane proteins with membrane-specific functions.

In summary, we present a protein design-driven approach to decipher mechanisms by which CDL regulates membrane protein stability. Our findings illuminate critical protein features that are challenging to predict or design *de novo*. Further integration of MD simulations and nMS with protein engineering could help to overcome some of the challenges identified here, such as the impact of local protein flexibility, and offer a promising pathway to uncover novel CDL binding sites. The findings not only contribute to our understanding of lipid-protein interactions but highlight potential avenues for the design of membrane proteins with tailored stability and function, potentially informing therapeutic strategies targeting membrane protein dysfunctions.

## Acknowledgments

The computations were enabled by resources provided by the National Academic Infrastructure for Supercomputing in Sweden (NAISS) at the PDC Center for High Performance Computing, KTH Royal Institute of Technology, partially funded by the Swedish Research Council through grant agreement (2022-06725). Additional computational resource came from ARCHER and JADE UK National Supercomputing Services, provided by HECBioSim, the UK High End Computing Consortium for Biomolecular Simulation (https://www.hecbiosim.ac.uk), which is supported by the EPSRC (EP/L000253/1). EM data was collected at the Karolinska Institutet 3D-EM facility (https://ki.se/cmb/3d-em). M.L.A is supported by a VR Research Environment grant (2019-02433). M.L. is supported by a KI faculty-funded Career Position, a Cancerfonden Project grant (22-2023 Pj), a VR Starting Grant (2019-01961), and a Consolidator Grant from the Swedish Society for Medical Research (SSMF). K.S. and J.S. have been supported by the Operational Programme European Regional Development Fund (no. CZ.02.1.01/0.0/0.0/16_019/0000729). K.S. also acknowledges support from the InterCOST programme of the Ministry of Education, Youth and Sports of the Czech Republic (project no. LUC23180).C.V.R. and A.O.O. are supported by the Medical Research Council (MRC) programme grant (MR/V028839/1). D.D. acknowledges support from Göran Gustafssons Foundation. O.A. was supported by a doctoral fellowship from Sven och Lilly Lawskis stiftelse. R.J.H. and E.L. by grants from the Swedish Research Council (2019-02433, 2021-05806) and Swedish e-Science Research Center. L.J.P and E.G.M are supported by a project grant from the Swedish Research Council (2020-04825). P.J.S. acknowledges Wellcome (208361/Z/17/Z), MRC (MR/S009213/1), BBSRC (BB/P01948X/1, BB/R002517/1, BB/S003339/1 and BB/Y003187/1), and the Howard Dalton Centre for funding. P.J.S. acknowledges Sulis at HPC Midlands+, which was funded by the EPSRC on grant EP/T022108/1, and the University of Warwick Scientific Computing Research Technology Platform for computational access.

## Data and materials availability

Cryo-EM density maps of ROCKET and ROCKET^AAXWA^ in detergent micelles have been deposited in the Electron Microscopy Data Bank under accession number EMD-50106 and EMD-50107 respectively. All data are available in the main text or the supplementary materials.

