## Supplementary material for "Engineering cardiolipin binding to an artificial membrane protein reveals determinants for lipid-mediated stabilization": (Fig S2)

**This PDF file includes:**

Materials and Methods
Figures S1 to S8
Tables S1 to S3
Legends for Movies S1
References

**Other supporting materials for this manuscript include the following:**

Movies S1

### Materials and Methods

Coarse-grained molecular dynamics simulations of ROCKET variants and GlpG.

CG systems were built using PDB 6B85 (TMHC4_R) or 2IC8/2NRF (GlpG). For the 6B85 system, two subunits of the heterotetramer were unchanged from the input PDB, whereas the other two were mutated to create the ROCKETAAXWA variant These mutations were added in PyMOL (*1*). Protein atoms were converted to the CG Martini 3 force field (*2*) using the martinize method (*3*). Additional bonds of 500 kJ mol−1 nm−2 were applied between all protein backbone beads within 1 nm. Proteins were built into membranes composed of 10% CDL, 23% POPG, and 67% POPE for ROCKET, or 10% CDL, 10% POPG, and 80% POPE or just 10% POPG and 90% POPE for GlpG. The default CDL Martini 3 parameters were used, corresponding to di-PO tail types. Membranes were built using the insane protocol (*4*). All systems were solvated with Martini waters and Na+ and Cl− ions to a neutral charge and 150 mM. Systems were minimized using the steepest descents method, followed by 1 ns equilibration with 5 fs time steps, then by 100 ns equilibration with 20 fs time steps, before 5 × 10 μs production simulations using 20 fs time steps, all in the NPT ensemble at 323 K with the V-rescale thermostat (τt = 1.0 ps) and semi-isotropic Parrinello-Rahman pressure coupling at 1 bar (τp = 12.0 ps). The reaction-field method was used to model long-range electrostatic interactions. Bond lengths were constrained to the equilibrium values using the LINCS algorithm. Density analyses was performed using the VolMap tool of VMD, with the default settings (*5*). Lipid binding sites and lipid-residue interactions were determined using the PyLipID package, which provides both occupancy and residence time data (*6*). Reported occupancy and residence time values are taken analysis of the total simulation time of 5 x 10 μs. Simulations were run in Gromacs 2022 (*7*, *8*).

Atomistic molecular dynamics simulations of ROCKET.

A post-CG simulation snapshot of ROCKET with bound CDL lipids was converted to an atomistic description using the CG2AT approach **(*9*)** Atoms were described using the CHARMM36m force field **(*10*)** with TIP3P water. Converted systems were energy minimized using the steepest descents method, and subsequently equilibrated with positional restraints on heavy atoms for 100 ps in the NPT ensemble at 303 K with the V-rescale thermostat (τp = 1.0 ps) and semi-isotropic Parrinello-Rahman pressure coupling at 1 bar (τp = 5.0 ps), with a compressibility of 4.5 × 10−5 bar−1. The Particle-Mesh-Ewald (PME) method was used to model long-range electrostatic interactions. van der Waals (VDW) interactions were cut off at 1.2 nm. Bond lengths were constrained to the equilibrium values using the LINCS algorithm. A production simulation was run to assess the dynamics of the bound lipid, using a 2 fs time step for 530 ns. Simulations were run in Gromacs 2022 **(*7*, *8*)**. A video was made using VMD.

Gas-phase molecular dynamics simulations of ROCKET variants.

In order to assess how lipid binding affects the stability of ROCKET and the ROCKET$^{AAWXA}$ variant in the gas-phase, molecular dynamics simulations were performed, in which chains were pulled apart and the force was measured with and without CDL bridging them. The proteins were simulated as homotetramers, using snapshots from the CG simulations which were converted into all-atom models. The chains were truncated to include only the membrane-spanning domain with residues S2-V79, with a neutral C-terminus (COOH) added to residue 79. In aiming to reflect the 16+ charge state, which was observed in mass spectrometry experiments, residues D38 and E78 were protonated, while all other titratable residues were given their pKa-based protonation state at pH 7. The addition of eight protons to the truncated model is intended to replicate the distribution of exposed acidic sites on the entire protein, which are likely to become protonated in the experiments. Each variant was simulated without any lipids, as well as with one CDL molecule bound between two helices of adjacent subunits.

Simulations were performed with the GROMACS MD package (*11*), version 2023.3, and the July 2022 version of the CHARMM36 force field (*12*). The proteins were placed in cubic boxes with sides of 999.9 nm, and the cutoff radii for Coulomb and van der Waals interaction were set to 333.3 nm. This set-up allows the use of the Verlet buffer scheme (*13*) for neighbour searching and GPU acceleration while avoiding artefacts from the periodicity (*14*). Virtual sites (*15*) were used for hydrogens, for which cardiolipin was generated with MkVsites (*16*). The four models were equilibrated with a steepest descent energy minimization until convergence, followed by temperature coupling over 10 ps, using a 0.5-fs time step, and the Berendsen (*17*) thermostat set to 300 K. For the production simulations, the thermostat was changed to velocity rescale

(*18*), retaining the temperature of 300 K. All bonds were constrained with LINCS(*19*), using an order of 4 and one iteration, which allowed for a 5-fs time step to be used.

Helices were pulled apart with the center-of-mass pull code. The reference groups included, respectively, the Cα atoms of residues 6-12 and 63-69 on the two chains with which CDL was interacting. The pull coordinate was defined as the distance between the mass centra of the two groups, measured in three-dimensional space. An umbrella potential with a harmonic force constant of 1000 kJ/mol/nm$^2$ and a pull rate of 0.1125 nm/ns were used. Twenty replicate pulling simulations of 50 ns each were performed for each of four models using a random seed for the velocities. The helix content was computed with the DSSP*(20) module of MDAnalysis (21, 22).*

Protein engineering of ROCKET mutants.

All protein mutagenesis was performed with the Q5 site-directed mutagenesis kit from NEB (E0554). Primers were designed with the NEBaseChanger tool (https://nebasechanger.neb.com/) and synthesized (Eurofins Genomics) with the sequences listed (Table S2). The original plasmid used as template for the PCR reactions was based on wildtype ROCKET plasmid (Table S3) and the reactions utilized the Q5 Hot Start High-Fidelity DNA Polymerases with annealing temperature optimized for each individual reaction (Table S2). PCR products were prepared using kinase, ligase, and DpnI (KDL) mixture, according to the manufacturer protocol, transformed into NEB 5-alfa cells, plated on Luria-Bertani (LB) agar plates containing kanamycin (50 µg/mL) and incubated overnight at 37°C. Individual colonies were selected for growth in 5 mL LB cultures at 37°C for 12-16 hours. 1-5 mL of cell pellet was harvested for plasmid extraction with the Monarch Plasmid DNA Miniprep Kit (New England Biolabs) (T1010). Mutagenesis was confirmed by sequencing.

Cloning and expression of ROCKET variants.

Synthetic genes with N-terminal histidine tag (either 6-His or 10-His for ROCKET$^{AAXWA}$) were synthesized by Genscript Inc or derived from mutagenesis in a pet26b (+) expression vector (Table S3). These plasmids were transformed into *E. coli* BL21 (DE3) (New England BioLabs). Selection was carried out on LB agar plates containing kanamycin (50 µg/mL) and incubated overnight at 37°C. Pre-cultures were grown under the same condition overnight and used to inoculate 400 mL LB media at a 1:100 dilution for protein expression. The expression cultures were incubated at 37°C until $OD_{600}$ reached 0.8-1.0, at which protein expression was induced with 0.2 mM isopropylthio-β-galactoside (IPTG). Post-induction, the cultures were incubated at 18°C overnight for protein expression. Cells were harvested by centrifugation at 8 000 x g for 20 minutes.

Purification of ROCKET variants.

Cell pellets were homogenized in resuspension buffer (25 mM TRIS pH 8.0, 150 mM NaCl) and subjected to probe sonication. n-Dodecyl-beta-maltoside (DDM) was added to the lysate to a final concentration of 1 % (w/v) and the mixture was incubated overnight at 4°C with shaking. Following solubilization, the solution was centrifuged at 10 000 x g for 10 minutes the supernatant was then filtered through a 0.2 µm syringe filter. The cleared supernatant was applied to a $Ni^{2+}$ Sepharose High Performance column HisTrap (Cytiva) pre-equilibrated with wash buffer (25 mM TRIS pH 8.0, 150 mM NaCl, 30 mM imidazole, and 0.1 % DDM). The column was subsequently washed with the same buffer to remove unbound material.

Protein was eluted with an elution buffer (25 mM TRIS pH 8.0, 150 mM NaCl, 300 mM imidazole and 0.1 % (w/v) DDM). Elution fractions were collected and analyzed for protein content by SDS-PAGE. Fractions containing the protein of interest were pooled and concentrated using Amicon Ultra-15 centrifugal filter units with a 100-kDa molecular weight cutoff (Merck Millipore) Concurrently, the buffer was changed to remove imidazole, resulting in a final storage buffer (25 mM TRIS pH 8.0, 150 mM NaCl and 0.1 % (w/v) DDM) for downstream applications.

Cryo-EM sample preparation and data acquisition.

5 mg/mL ROCKET and ROCKET$^{AAXWA}$ with 100 µM CDL were frozen on Quantifoil 1.2/1.3 Au 300 mesh grids (Quantifoil Micro Tools). 3 µL of sample was applied to each grid which was then blotted for 3 s and plunge-frozen into liquid ethane using FEI Virtobot Mark IV (Thermo Fisher Scientific). Micrographs were

collected on Krios G3i electron microscope (Thermo Fisher Scientific) operated at 300 kV equipped with Gatan BioQuantum K3 image filter and a Ceta-D detector. Movies were collected at a nominal 165,000 x magnification, resulting in a pixel size of 0.5076 Å. A total dose of 60 $e^{-}/Å^{2}$ was used to collect 29 frames over 1 s. The target defocus range was set between -0.6 to -1.8 µm, in steps of 0.2 µm.

Image processing & model building.

Data processing was performed using the RELION 4.0.1 pipeline (*23*). Motion correction was performed using Relion's own implementation (*24*) and CTF estimation was done with CtfFind4.1 (*25*). For the WT dataset, 2D references from 2D classification of manually picked particles were used for initial autopicking, and the picked particles were used to generate a low-resolution 3D reconstruction that was used and a 3D reference for the final round of autopicking. Due to high similarity in 2D classes from manually picked particles for the WT and MUT5 dataset, the same low resolution 3D reconstruction from the WT processing pipeline was used directly as a 3D reference for autopicking in the MUT5 data set. For both datasets, the auto picked particles were used for an ab initio reconstruction. The particles were then further refined, and the data cleaned using several rounds of 3D classification and 3D auto-refinement, followed by CTF parameters refinement and particle polishing before a final 3D auto-refinement and post-processing. No smaller particles were identified during manual and automated processing.

Preparation of proteoliposomes for native mass spectrometry

*E. coli* polar lipid extract, with a composition of 67.0% PE, 23.2% PG, 9,8% CDL (Avanti Polar Lipids) was dissolved in a 1:1 mixture of chloroform and methanol. The solvent was then removed under vacuum using a SpeedVac concentrator (Savant SPD1010) until completely dry. The dried lipid film was resuspended in buffer (25 mM TRIS pH 8.0, 150 mM NaCl) and vortexed vigorously to ensure homogeneity. Large unilamellar vesicles (LUVs) were prepared by extrusion though a pair of 0.4 µm polycarbonate membrane. The target protein-to-lipid ration was 1:100 (protein:lipid by weight) and the mixture was incubated at 37°C for 30 minutes to allow for protein incorporation into the lipid vesicles. Post-incubation, the samples were dialyzed against 500 mM ammonium acetate, pH 8.0, overnight to facilitate buffer exchange and removal of remaining detergent. The particle size distribution of LUVs and proteoliposomes were monitored by dynamic light scatter (DLS) using a Viscotek model 802 DLS instrument with an internal laser (825-832 nm). Data processing was performed with OmniSIZE2.

Native mass spectrometry of proteoliposomes.

Proteoliposome samples were subjected to ESI nMS were performed on the Q Exactive Ultra-high range (UHMR) mass spectrometer (Thermo Fisher Scientific). The MS capillaries were custom-pulled and coated in-house (*26*). We set the capillary voltage to 1.5 kV and maintained the source temperature at 270°C. In-source trapping voltage was applied at 300 V to enhance ion desolvation, and the higher-energy collision dissociation (HCD) voltage was set to 200 V to facilitate ion transmission. The ultra-high vacuum pressure within the MS was measured at $6.01 \times 10^{-10}$ mbar. Data was analyzed using Xcalibur 2.2 (Thermo Fisher).

Protein preparation for native mass spectrometry.

Immediately prior to MS analysis, the purified protein was subjected to size exclusion chromatography (SEC) using a Superdex 200 Increase 10/300 GL column (Cytiva). The detergent exchange process was conducted with native compatible buffer (200 mM ammonium acetate pH 8.0, 0.5% C8E4). The fractions containing the tetrameric state of ROCKET protein was collected for direct analysis with nMS.

Native mass spectrometry.

ESI-MS spectra were recorded on a Waters Synapt G1 wave ion mobility mass spectrometer, modified for high-mass analysis (MS Vision), and equipped with an offline nanospray source. The ESI-MS parameters were set as follows: capillary voltage at 1.5 kV, cone voltage at 100 V, source pressure maintained at 8 mbar, and source temperature regulated at 30°C. To optimize the detection of protein-detergent complexes, the trap voltage was varied from 90-240 V. For assessments of lipid binding ROCKET and $ROCKET^{AAXWA}$ variants were prepared in samples with and without 50 µM 16:0 cardiolipin (Avanti Polar Lipid). The nMS setting were the same as above, with a trap voltage of 170 V. For each condition, four protein-lipid mixtures were prepared and measured separately. The mass spectra were analyzed using MassLynx software version 4.1 (Waters), and the intensities of apo- and lipid-bound tetramers were quantified with mMass V3.9.0.

To measure lipid-mediated stabilization, 15 µM ROCKET or ROCKET$^{MUT}$ were mixed with 15 µM ROCKET$^{AAXWA}$ at an equimolar ratio to achieve comparable intensities for tetrameric species at a collision voltage of 170 V. The mixtures were the supplemented with 16:0 cardiolipin (Avanti Polar Lipid) to a final lipid concentration of 25 µM. nMS setting were consistent with previous experiments and the collision voltage set to 200-220 V to allow optimal detection of unfolded monomers. Four spectra were recorded for each condition (n=4). The mass spectra were analyzed using MassLynx software version 4.1 (Waters), and the intensities of the charge state of the monomers were quantified with mMass V3.9.0.

Screening of lipid impact on of GlpG-GFP fusion protein stability.

GlpG-GFP fusion protein was expressed and purified as described previously (*27*). The purified fusion protein was diluted in buffer containing 20 mM Tris-HCl pH 8.0, 150 mM NaCl, 1 % (w/v) β-OG and 1% (w/v) DDM to a final concentration of fusion protein of 1 µM, and individual lipids were added to this preparation to a final concentration of 0.3 mg/mL. The used lipids were from Avanti Polar Lipids: 18:1 PE (cat no. 850725P), 18:1 PG (cat no. 840475P), 18:1 PC (cat no. 850375P), 18:1 CDL (cat no. 710335P) and brain SM (cat no. 860062P), and DDM was used as a negative control. Samples were incubated at 63°C for 10 min (the negative control at 4°C) followed by centrifugation at 20 000xg at 4°C for 45 minutes. Fluorescence of the supernatant was measured with TECAN Infinite M1000 spectrophotometer with excitation at 488 nm and emission at 512 nm.

Determination of protease activity of GlpG.

To determine GlpG activity, the GlpG-GFP supernatant collected after centrifugation was diluted 1:10 into 50 mM phosphate buffer pH 7.4, 150 mM NaCl, 0.05 % (w/v) PEG 8000, 20 % (v/v) glycerol, and 0.05 % (w/v) DDM. Lyophilized substrate were dissolved in the same buffer, with further addition of 5% (v/v) DMSO in case of the soluble substrate KSp63 (*28*), and preincubated at 37°C. The concentration of substrates in these master mixes were 400 µM for the ‘soluble’ (extramembrane) substrate KSp63 and 50 µM for the transmembrane substrate KSp96 (*29*). Cleavage reactions were initiated by mixing the enzyme solution and substrate master mix in 1:1 ratio, yielding final concentrations of 200 µM KSp63 and 25 µM KSp96. Protease activity was measured at 37°C by reading fluorescence intensity continuously every 30 seconds in TECAN Infinite M1000 for 40 min or until the upper detection limit was reached. Excitation was set to 355 nm and emission to 450 nm for the soluble substrate KSp63, or to 335 nm and 493 nm, respectively, for the transmembrane substrate KSp96.

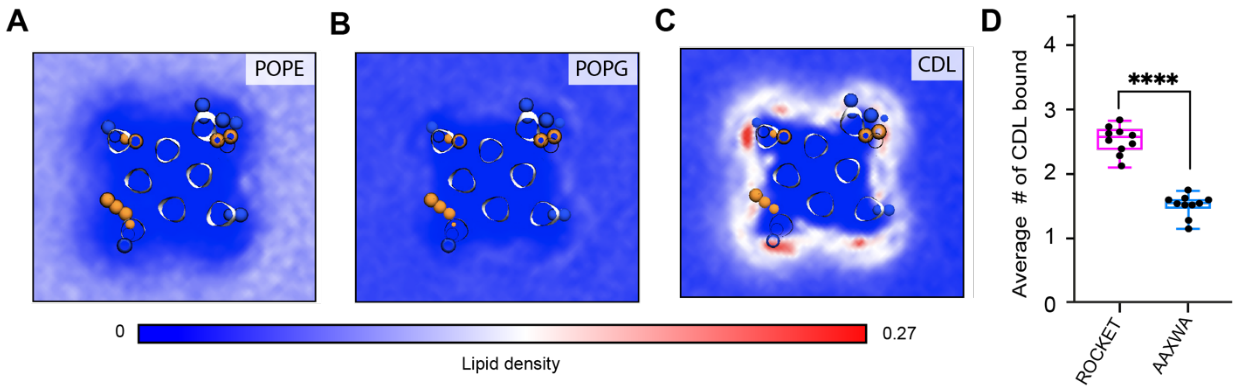


**Fig. S1.**

**CG-MD simulations of lipid interactions with ROCKET and ROCKET$^{AAXWA}$.**

CG-MD-derived lipid densities around tetrameric ROCKET for all lipid components in the model *E. coli* membrane, *(A)* POPE, *(B)* POPG, and *(C)* CDL. Only CDL displays preferential binding. POPE; on the other hand, is largely excluded from the protein surface.

*(D)* The average number of CLD molecules bound to tetrameric ROCKET and ROCKET$^{AAXWA}$ in CG-MD simulations, as analyzed using gmx select with a distance of 0.6 nm between CDL headgroup and the protein surface used as cut-off, see Methods for details. Statistics are from two-tailed t-tests (n=5) (**** p<0.0001).

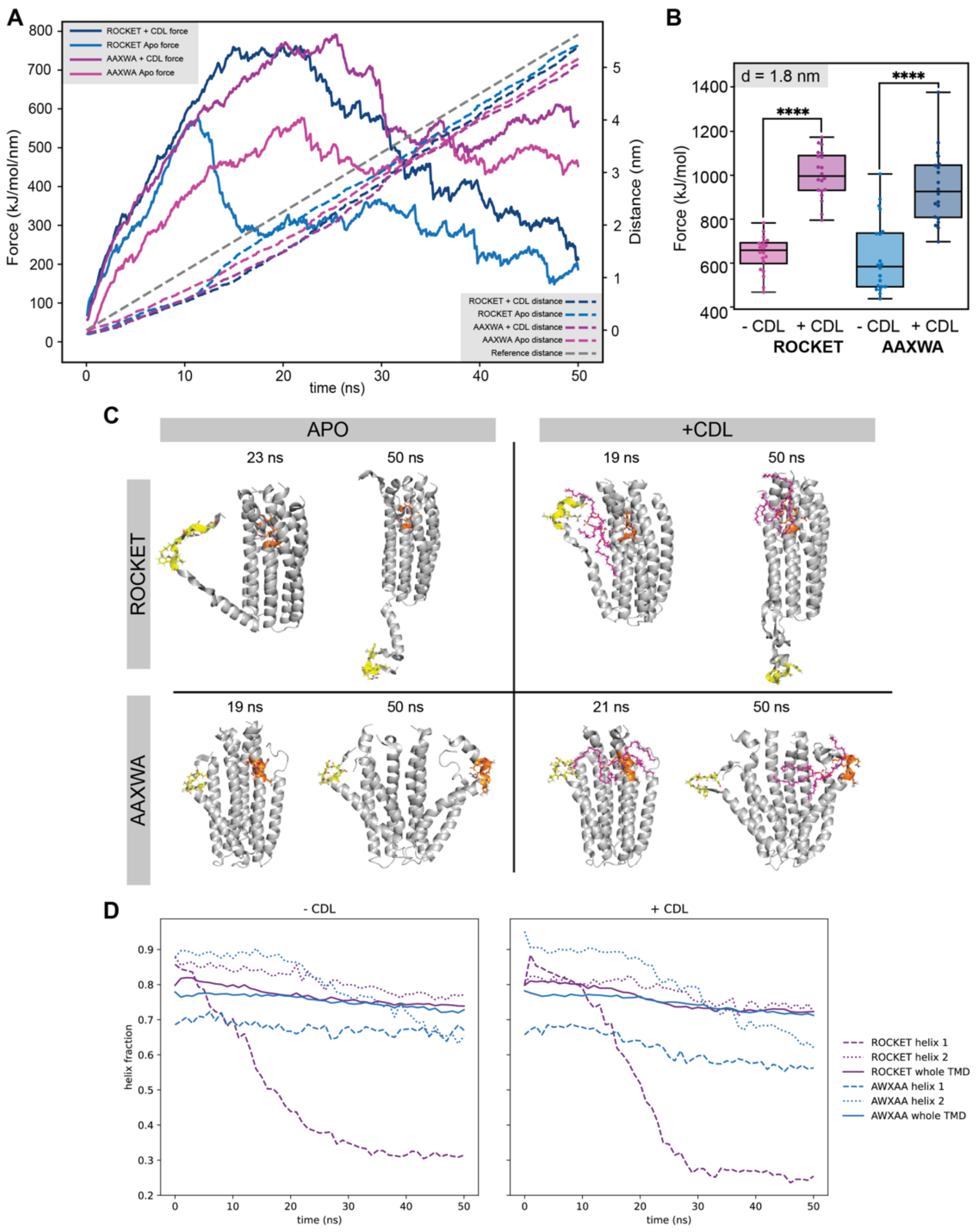


**Fig. S2.**
**Molecular dynamics simulations of ROCKET in the gas phase**.
*(A)* Trajectory of gas phase simulations showing the force (dashed lines) and the total distance between the N- and C-terminal ends of helices 1 and 2, respectively (solid lines) over the 50 ns simulation for ROCKET and ROCKET$^{AAXWA}$ with and without CDL. The measurements are averages of the 20 replicate simulations. At around 10 ns, the contacts between helix 1 and 2 are disrupted.
*(B)* Plots of the total force required to separate helix 1 and 2 by d = 1.8 nm for ROCKET (purple) (p =1.99*10$^{-14}$) and ROCKET$^{AAXWA}$ (blue) (p = 5.34*10$^{-7}$) with and without bound CDL. At 1.8 nm, the core of the tetramer dissociates, which is prevented by CDL for both ROCKET variants (two-tailed t-test, n=20).
*(C)* Snapshots of unfolding trajectory of ROCKET and ROCKET$^{AAXWA}$ in the absence (left) and presence (right) of CDL at 19-20 ns, where helix 1 is completely dissociated, and at the end of the simulations (50 ns).
*(D)* Helix content of ROCKET and ROCKET$^{AAXWA}$ over the course of the gas phase unfolding simulations. The conformation of Helix 1 in ROCKET is stabilized by CDL binding. The AAXWA mutation stabilizes the helical conformation independently of CDL binding.

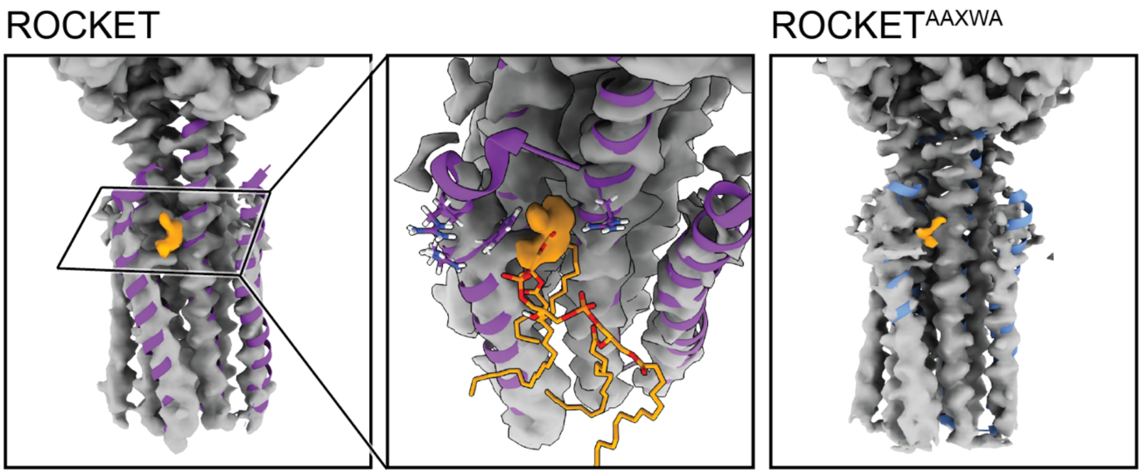


**Fig S3.**
**Cryo-EM densities for ROCKET and ROCKET$^{AAXWA}$ in the presence of CDL**.
The MD-relaxed backbone of the X-ray structure of ROCKET (shown as ribbons) largely aligns with the densities. No substantial differences can be observed between the densities determined for ROCKET (left panel) and ROCKET$^{AAXWA}$ (right panel). The N-terminal end of helix 1 (residues 1-12) is relatively poorly resolved, possibly due to intrinsic flexibility. For both variants, a diffuse non-protein density (orange) partially overlaps with the lipid headgroup of CDL predicted in site 2 (middle panel).

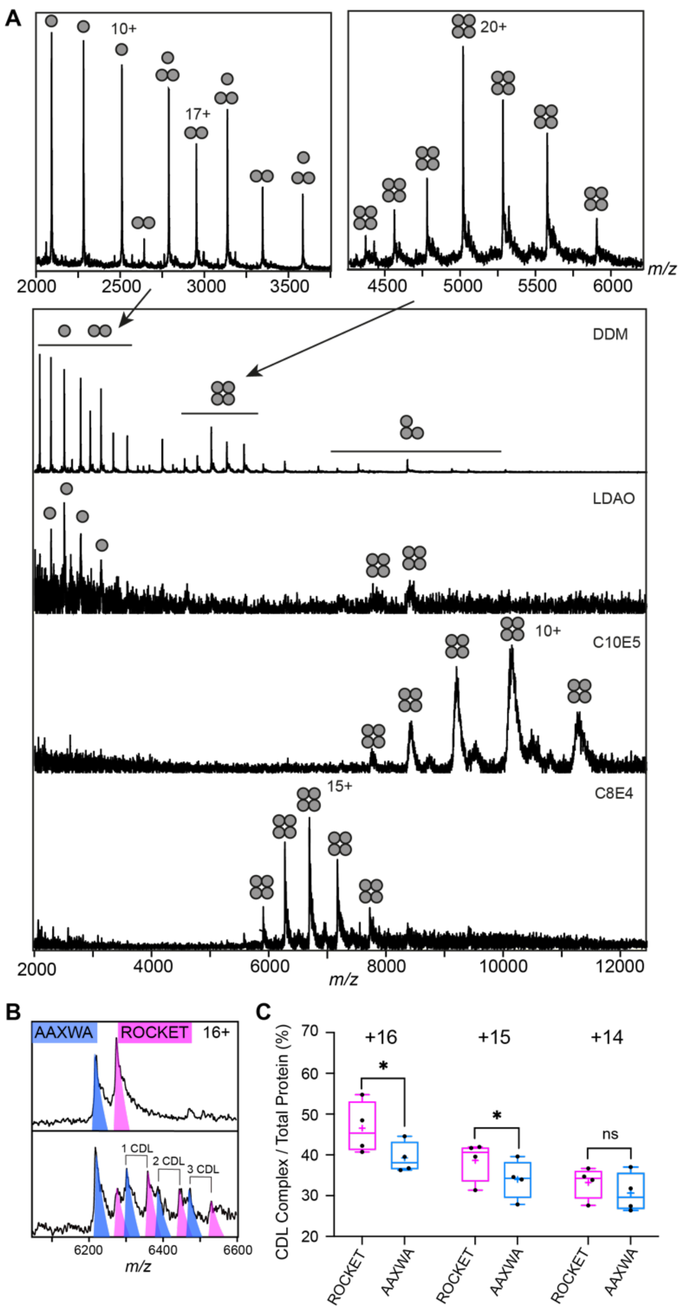


**Fig S4.**

**Detergent screening and CDL binding to ROCKET and ROCKET$^{AAXWA}$.**

*(A)* Detergent screening of ROCKET with nMS. Detergents DDM, LDAO, C10E5, and C8E4 were compared, only the PEG detergent C8E4 yielded intact tetramers.

*(B)* CDL binding assay with equimolar amounts of ROCKET and ROCKET$^{AAXWA}$ in C8E4 before (top) and after (below) addition of CDL show preferential binding to ROCKET. The 16+ charge state is shown. Spectra were recorded at collision voltage of 170V.

*(C)* Quantification of lipid binding difference between ROCKET and ROCKET$^{AAXWA}$ (as shown in Figure S4b) for the main charge states. The higher charge states (+16 and +15) show a significant change in lipid binding between ROCKET and ROCKET$^{AAXWA}$, indicating that positively charged residues contribute to CDL binding (two-tailed t-tests (n = 4), p = 0.0140 and p = 0.0284 for 16+ and 15+, respectively, p = 0.1733 for 14+).

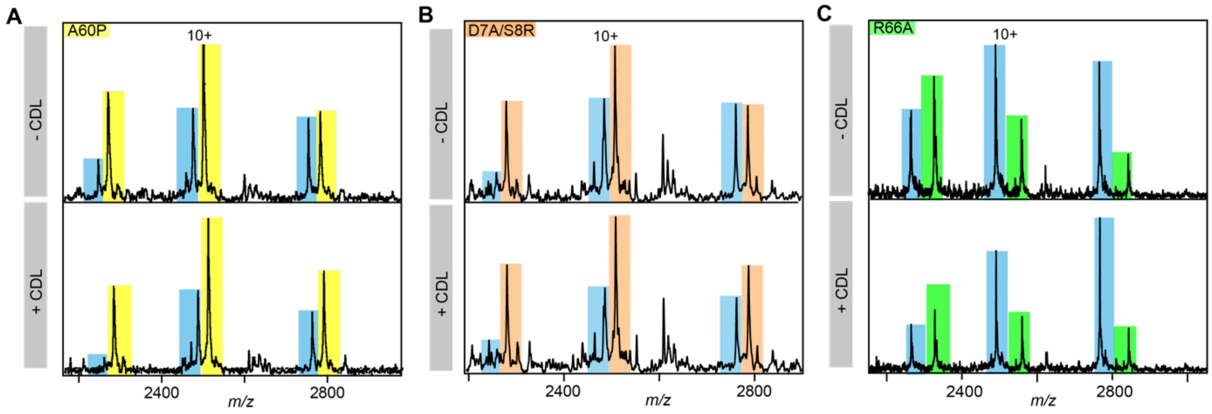


**Fig S5.**
**nMS spectra showing lipid-mediated stabilization** of ROCKET$^{AAXWA}$ and ROCKET$^{R66A}$ (yellow). Representative nMS spectra showing the intensity changes for unfolded monomers released from ROCKET$^{AAWXA}$ (blue) and *(A)* ROCKET$^{A60P}$ (yellow), *(B)* ROCKET$^{D7A/S8R}$ (orange), and *(C)* ROCKET$^{R66A}$ (green). See Figure 3 for quantification.

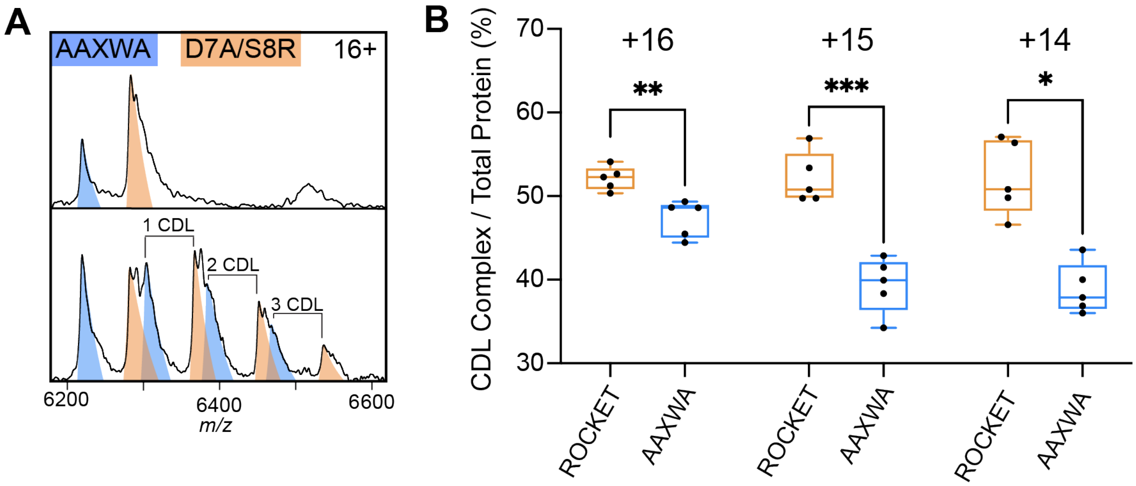


**Fig S6.** *(A)* CDL binding assay with equimolar amounts of ROCKET$^{D7A/S8R}$ and ROCKET$^{AAXWA}$ in C8E4 before (top) and after (below) addition of CDL show preferential binding to ROCKET. The 16+ charge state is shown. Spectra were recorded at collision voltage of 170V.

*(B)* Quantification of lipid binding difference between ROCKET$^{D7A/S8R}$ and ROCKET$^{AAXWA}$ (as shown in Figure S6a) for the main charge states. The higher charge states (+16 and +15) show a significant change in lipid binding between ROCKET$^{D7A/S8R}$ and ROCKET$^{AAXWA}$, indicating that positively charged residues contribute to CDL binding (two-tailed t-tests (n = 5), p = 0.0039 and p = 0.0009 for 16+ and 15+, respectively, p = 0.0102 for 14+).

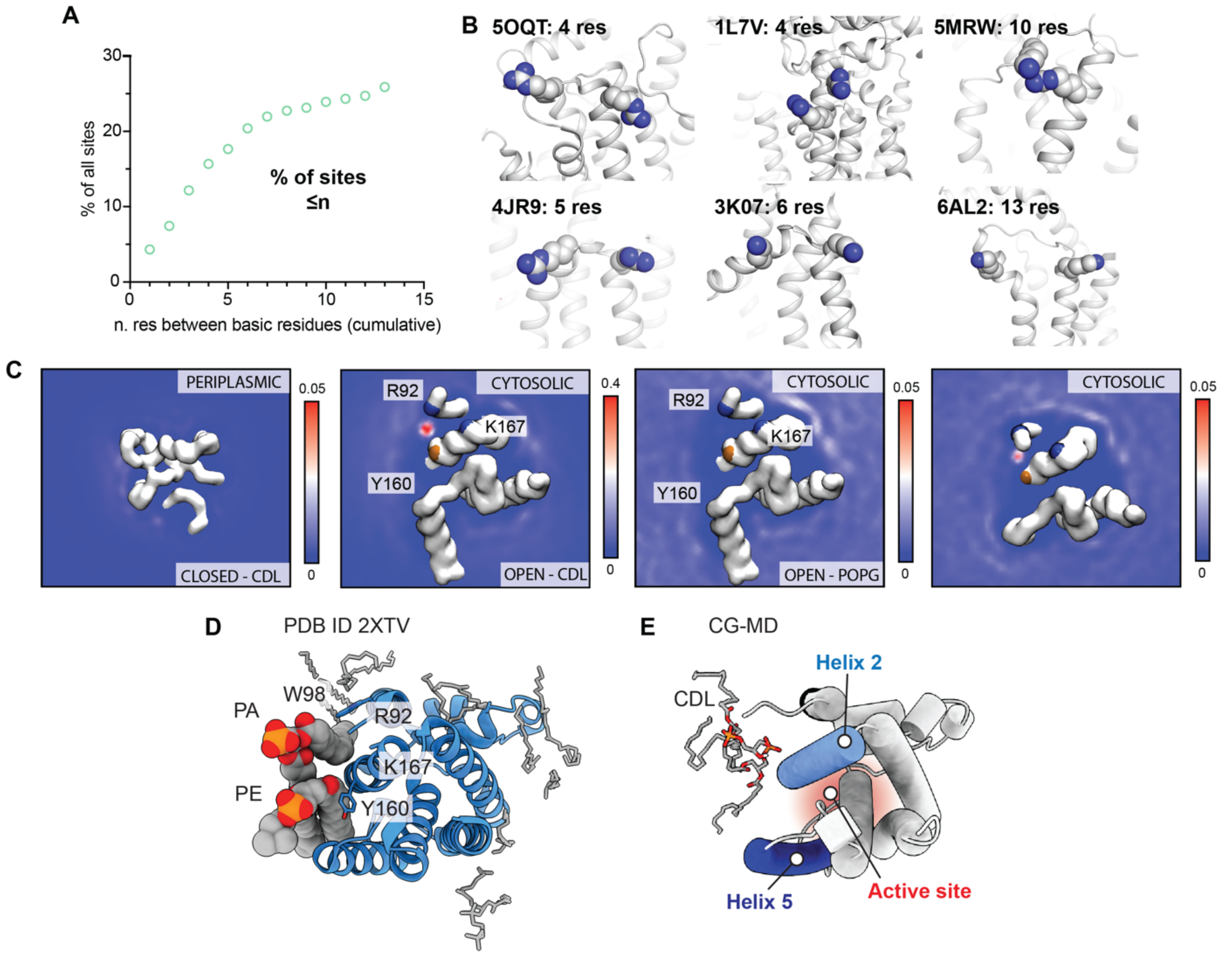


**Fig S7.**

**Analysis of CDL binding sites in *E. coli* proteins and GlpG**

*(A)* Distributions of distances between basic residues within single CDL binding sites in *E. coli* membrane proteins. Less than 20% of all binding sites contain basic residues separated by < 10 sequence positions, which is a requirement for CDL binding to a single helix.

*(B)* Representative CDL binding sites with less than ten residues between the basic residues show confinement to a single helix or loop region. Sites were identified by CG-MD simulations, see Ref. 12.

*(C)* Lipid density maps derived from CG-MD simulations of the closed state (PDB ID 2NRF) and open state (PDB ID 2IC8) of GlpG. The left panel shows the periplasmic side, the other panels the cytoplasmic side as in Figure 4d. The ‘CDL’ data are from a 10% CDL, 10% POPG, and 80% POPE membrane, whereas the ‘POPG’ data are from a 10% POPG and 90% POPE membrane. All density plots are taken over the total 5 x 10 µs of CG-MD.

*(D)* Crystal structure of GlpG with bound lipids shows tentatively modeled PA and PE lipids occupying the CDL binding site (PDB ID 2XTV).

*(E)* CG-MD snapshot of CDL bound to the closed conformation of GlpG, showing insertion of acyl chains into the gap between helices 2 and 5 that provides access to the active site on the other leaflet (red).

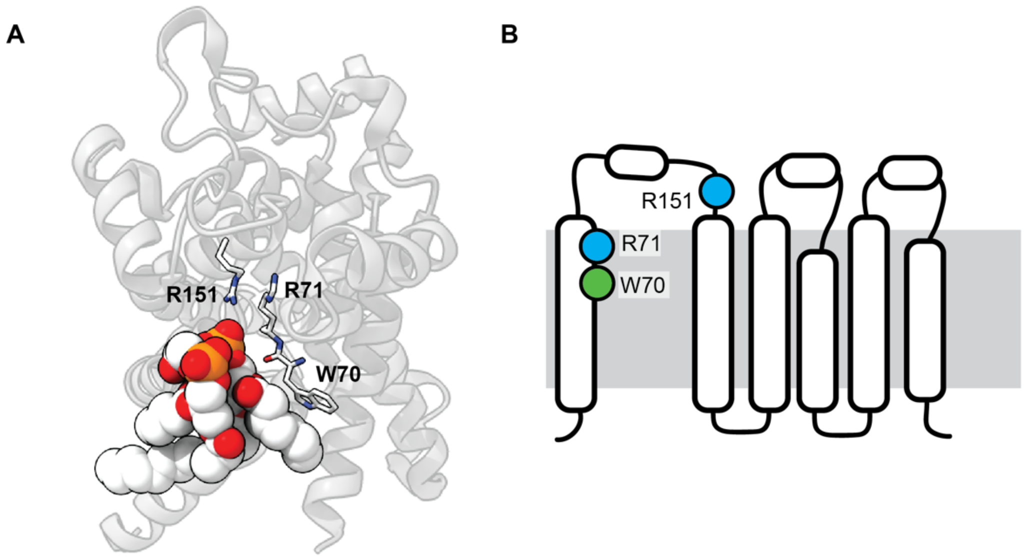


**Fig S8.**
**Analysis of the stabilizing CDL binding site in the Yeast ADP/ATP carrier Aac2**
*(A)* Crystal structure of Aac2 shows a CDL bound to W70/R71/R151. The residues coordinating CDL are shown as sticks, the CDL molecule is rendered as spacefill (PDB ID 1OKC). *(B)* The CDL site connects helix 1 to the helical core of Aac2, with high overall similarlty to the CDL site in GlpG.

**Table S1.**
Cryo-EM data collection and processing statistics.

| Data collection and processing | ROCKET | ROCKET$^{AAXWA}$ |
|---|---|---|
| Magnification | 165,000 | 165,000 |
| Voltage (kV) | 300 | 300 |
| Electron exposure (e$^{-}$/Å$^{2}$) | 60 | 60 |
| Defocus range (μm) | -0.6 to -1.8 | -0.6 to -1.8 |
| Pixel size (Å) | 0.5076 | 0.5076 |
| Number of images | 10,002 | 10,000 |
| Symmetry imposed | C4 | C4 |
| Particles refined | 98,241 | 142,434 |
| Map resolution (Å)<br>FSC threshold | 3.77<br>0.143 | 3.89<br>0.143 |
| Map sharpening B-factor | -133 | -164 |

**Table S2.**
Forward and reverse primers for mutagenesis based on the ROCKET sequence.

| Mutation | Primer sequence (**Forward**/<u>Reverse</u>) | $T_m$ (C°) |
|---|---|---|
| R9A/K10A/R13A (AAXWA) | **ctgggcgACCATTATGCTGTTACTG**<br><u>GTTTTCTGTGTCTTCTATCAcgccgcta</u> | 57 |
| A61P | **GGTTGTTATTccgCTGTTGCTCAG**<br><u>CGAATAGCGGAATTACGA</u> | 60 |
| D7A/S8R | **tcgcAAGATCTGGCGTACCATTATG**<br><u>CTATATGTATACAGTTTTCTGTGTCTTcgagc</u> | 62 |

**Table S3.**
Protein variants; expressed sequence and theoretical molecular weight (MW).

| Protein | Protein sequence (<u>Purification tag)</u> | Theoretical MW (Da) |
|---|---|---|
| ROCKET | MSKDTEDSRKIWRTIMLLLVFAILLSAIIWYQITTNPDTSQIA<br>TLLSMQLLLIALMLVVIALLLSRQTEQVAESIRRDVSALAYV<br>MLGLLLSLLNRLSLAAEAYKKAIELDPNDALAWLLLGSVLE<br>KLKRLDEAAEAYKKAIELKPNDASAWKELGKVLEKLGRLD<br>EAAEAYKKAIELDPEDAEAWKELGKVLEKLGRLDEAAEAY<br>KKAIELDPNDLE<u>HHHHHH</u> | 25 226.44 |
| ROCKET$^{AAXWA}$ | MSKDTEDSAAIWATIMLLLVFAILLSAIIWYQITTNPDTSQIA<br>TLLSMQLLLIALMLVVIALLLSRQTEQVAESIRRDVSALAYV<br>MLGLLLSLLNRLSLAAEAYKKAIELDPNDALAWLLLGSVLE<br>KLKRLDEAAEAYKKAIELKPNDASAWKELGKVLEKLGRLD<br>EAAEAYKKAIELDPEDAEAWKELGKVLEKLGRLDEAAEAY<br>KKAIELDPNDLE<u>HHHHHH</u> | 24 999.13 |
| ROCKET$^{A61P}$ | MSKDTEDSRKIWRTIMLLLVFAILLSAIIWYQITTNPDTSQIA<br>TLLSMQLLLIALMLVVIPLLLSRQTEQVAESIRRDVSALAYV<br>MLGLLLSLLNRLSLAAEAYKKAIELDPNDALAWLLLGSVLE<br>KLKRLDEAAEAYKKAIELKPNDASAWKELGKVLEKLGRLD<br>EAAEAYKKAIELDPEDAEAWKELGKVLEKLGRLDEAAEAY<br>KKAIELDPNDLE<u>HHHHHH</u> | 25 252.48 |
| ROCKET$^{D7A/S8R}$ | MSKDTEARRKIWRTIMLLLVFAILLSAIIWYQITTNPDTSQIA<br>TLLSMQLLLIALMLVVIALLLSRQTEQVAESIRRDVSALAYV<br>MLGLLLSLLNRLSLAAEAYKKAIELDPNDALAWLLLGSVLE<br>KLKRLDEAAEAYKKAIELKPNDASAWKELGKVLEKLGRLD<br>EAAEAYKKAIELDPEDAEAWKELGKVLEKLGRLDEAAEAY<br>KKAIELDPNDLE<u>HHHHHH</u> | 25 251.54 |
| ROCKET$^{R66A}$ | MSKDTEDSRKIWRTIMLLLVFAILLSAIIWYQITTNPDTSQIA<br>TLLSMQLLLIALMLVVIALLLSAQTEQVAESIRRDVSALAYV<br>MLGLLLSLLNRLSLAAEAYKKAIELDPNDALAWLLLGSVLE<br>KLKRLDEAAEAYKKAIELKPNDASAWKELGKVLEKLGRLD<br>EAAEAYKKAIELDPEDAEAWKELGKVLEKLGRLDEAAEAY<br>KKAIELDPNDLE<u>HHHHHHHHHH</u> | 25 689.90 |

**Movie S1 (separate file).**

View of Site 1 and 2 from ROCKET, with bound CDL. Arginine 65 (R65) of Site 2 is on the left in blue sticks, and arginine 9 (R9), lysine 10 (L10), and arginine 13 (R13) of Site 1 are on the right. Tryptophan 12 (W12), which contributes to both Site 1 and 2, is shown in the center in orange sticks. CDL molecules are shown in coloured sticks, and the protein in white cartoon.
